# Modeling the role of mortality-based response triggers on the effectiveness of African swine fever control strategies

**DOI:** 10.1101/2021.04.05.438400

**Authors:** Gustavo Machado, Trevor Farthing, Mathieu Andraud, Francisco Paulo Nunes Lopes, Cristina Lanzas

## Abstract

African swine fever (ASF) is considered the most impactful transboundary swine disease. In the absence of effective vaccines, control strategies are heavily dependent on mass depopulation and movement restrictions. Here we developed a nested multiscale model for the transmission of ASF, combining spatially explicit network model of animal movements with a deterministic compartmental model for the dynamics of two ASF strains within-pixels of 3 km x 3 km, amongst the pig population in one Brazilian state. The model outcomes are epidemic duration, number of secondary infected farms and pigs, and distance of ASF spread. The model also shows the spatial distribution of ASF epidemics. We analyzed quarantine-based control interventions in the context of mortality trigger thresholds for the deployment of control strategies.

The mean epidemic duration of a moderately virulent strain was 11.2 days assuming the first infection is detected (best-case scenario) and 15.9 days when detection is triggered at 10 % mortality. For a highly virulent strain, the epidemic duration was 6.5 days and 13.1 days, respectively. The distance from the source to infected locations and the spatial distribution was not dependent on strain virulence. Under the best-case scenario, we projected an average number of infected farms of 23.77 farms and 18.8 farms for the moderate and highly virulent strains, respectively. At 10% mortality-trigger, the predicted number of infected farms was on average 46.27 farms and 42.96 farms, respectively. We also demonstrated that the establishment of ring quarantine zones regardless of size (i.e., 5 km, 15 km) was outperformed by backward animal movement tracking. The proposed modeling framework provides an evaluation of ASF epidemic potential, providing a ranking of quarantine-based control strategies that could assist animal health authorities in planning the national preparedness and response plan.

## Introduction

African swine fever (ASF) has not been reported in South America since the 1980s (Costard et al., 2009), but it is widespread in the northern hemisphere, including most of Asia (Mighell and Ward, 2021), and stretched as far as Germany, where it continues to infect wild boars and domestic pigs throughout several countries (Gao et al., 2020; Sauter‐Louis et al., 2020). The rapid spread of ASF throughout the northern hemisphere (Yoon et al., 2020) has reignited concerns about ASF reintroduction into the Americas, especially with the recent detection in the Dominican Republic

The main modes of ASF transmission are direct contact with infected animals, ingestion of infected pork products and or residuals, and contact with contaminated fomites (Chenais et al., 2019). Overall, the between-farm pig transportation has been described as the major pathway of ASF propagation (Andraud et al., 2019; Chenais et al., 2019; Hayes et al., 2020; Liu et al., 2020). In addition, the swine population in ASF-free regions is immunologically naive as there are no effective vaccines (Dellicour et al., 2020; Dixon et al., 2020). All these factors make controlling the spread of ASF a very challenging problem. In European and North American countries, ASF response plans include national movement standstill, testing, and stamping out, pre-emptive depopulation of neighboring herds, enhanced surveillance in predefined control zones, contact tracing, and the enhancement of on-farm biosecurity (Gallardo et al., 2015; Bellini et al., 2016; Halasa et al., 2016a; Sánchez-Cordón et al., 2018; USDA-APHIS, 2020).

Evaluating the effectiveness and economic impact of control strategies (e.g., animal movement restrictions, mass depopulation) have been among the priorities for ASF-free countries like the US and Brazil (Halasa et al., 2018a; Hayes et al., 2020; USDA-APHIS, 2020). In this vein, disease spread simulation models have been widely used to study transmission dynamics and to evaluate control options (Bradhurst et al., 2015; Machado et al., 2020; Galvis, Prada et al., 2021), several examples of model outputs implementation are described (Heesterbeek et al., 2015; Ezanno et al., 2020; Galvis et al., 2020; Halasa, Græsbøll et al., 2020; Hayes et al., 2020). Disease detection based on pig mortality and clinical disease has been recommended as the most effective surveillance option for disease response to trigger subsequent investigations and stamping out of affected herds (Guinat et al., 2017; Nielsen et al., 2017; USDA-APHIS, 2020; European Commission, 2020). The effectiveness of using mortality data for early detection of ASF is supported by additional modeling studies (Guinat et al., 2018; Faverjon et al., 2020).

Even though mathematical models have been proposed to estimate epidemiological consequences of ASF introduction in European countries (Halasa et al., 2016a, 2018a; Andraud et al., 2019), there are gaps in such models mainly because known variations on ASF clinical presentation and mortality patterns were not considered. Indeed, ASF propagation is dependent on multiple factors, including modes of disease introduction and dose of viral infection, the virulence of the strain, as well as host and population factors (Salguero, 2020; Mighell and Ward, 2021). In this study, we developed a nested multiscale model for the transmission of two ASF strains, with distinct virulence characteristics. Transmission among pixels is represented by a spatially explicit network model of animal movements. The spatial unit was defined as 3 km x 3 km gridded cells (pixel), in which the within-pixel ASF dynamics are represented by a deterministic compartmental model. The model provides epidemic duration, secondary infections, and distance spread across all pig populations of one Brazilian state. We also investigated how early detection based on mortality thresholds affects future epidemic outcomes while producing maps of secondary cases. Finally, we compared and contrasted different control strategies, namely: established control zones, contact tracing, with different surveillance mortality-base trigger thresholds scenarios, and including simulating the spread of high and moderate virulence strains. We demonstrate the utility of spatially explicit models in producing maps to guide preparedness activities in ASF‐free regions. All results are presented in the context of mortality-trigger and virulence combinations.

## Material and Methods

In the sections below we describe i) data sources used to recreate the temporally-dynamic pig shipment network model; ii) the design and parameterization of the ASF transmission model simulated on the between-farm movement network; iii) the model inputs we used to simulate the spread of ASF strains, and iv) the model output and analyses employed to assess the disease control scenarios.

### Study areas and animal movement data

The study area comprises the swine population of Rio Grande do Sul, Brazil. The state holds nearly 15% of the national commercial pig population. More details about the pig population and a detailed description of the multiyear between farm pig movements networks are described elsewhere (Machado et al., 2020). Here, we reconstructed the between-farm pig transportation network describing animal shipments from Jan. 2^nd^, 2015 until Dec. 31^st^, 2018. This network describes a total of 273,158 shipments between 13,111 registered swine farms. For each individual pig farm, we identified: farm geolocation, for commercial operations pig company name to which the farm(s) are contracted with, pig capacity, and the number of pigs per premise, from the local official veterinary service (SEAPI-RS, 2018). Farms declared inactive, in which no pigs were raised between 2017 and 2018, or that were reported to be out of business, were not considered in the analysis. Movements from or to other states were also excluded, given the lack of information about farm registration outside the Rio Grando do Sul state lines.

The study area, the entire state of Rio Grande do Sul (Figure 1), was divided into gridded 3 km x 3 km cells (pixels) to which the between-farm pig movements were aggregated. Briefly, each farm location was intersected into one of 5,143 unique 3-km pixels containing at least one pig farm. The number of farms contained within pixels ranged from one to a maximum of 6,638, with a median count of two farms in each pixel. The 3-km pixel size was based on previous work (Halasa et al., 2016a), in which extensive sensitivity analyses suggested that a distance of up to 2 km around infected farms adequately capture the mixture of unregistered animal movements, shared equipment, and tools, people or sharing of equipment between neighbors, and local transmission events driven by rodents and insects and other wildlife hosts. Thus, by creating a pixel-level network, we were able to minimize computational requirements without sacrificing model realism. The pixel-level network consisted of 259,004 temporally-dynamic daily directed edges between 4,367 pixels. This was a network with no edge (i.e., animal movement) originating and ending at the same node (pixel), meaning that no within-pixel farm-to-farm animal shipments were represented. On average for the entire network, pixels had daily in- and out- degrees of 0.04 (see Supplementary Figure S1 for the description of daily movements within-pixel farm-to-farm and between-pixel total, in-, and out- degree). For modeling purposes, we calculated the summed pig-holding capacity of all farms within each pixel in the network and appended this information to the network as a pixel-level variable to be used as a proxy for swine population size. Estimated pixel-level swine population values in the network ranged from 0 to 86,834 pigs. When summed farm-level pig-holding capacities were unknown or reported to be 0, pixels were assigned capacities of 0. For each pixel, we also identified what swine production companies operated ≥ 1 farm contained therein.

**Figure 1.**
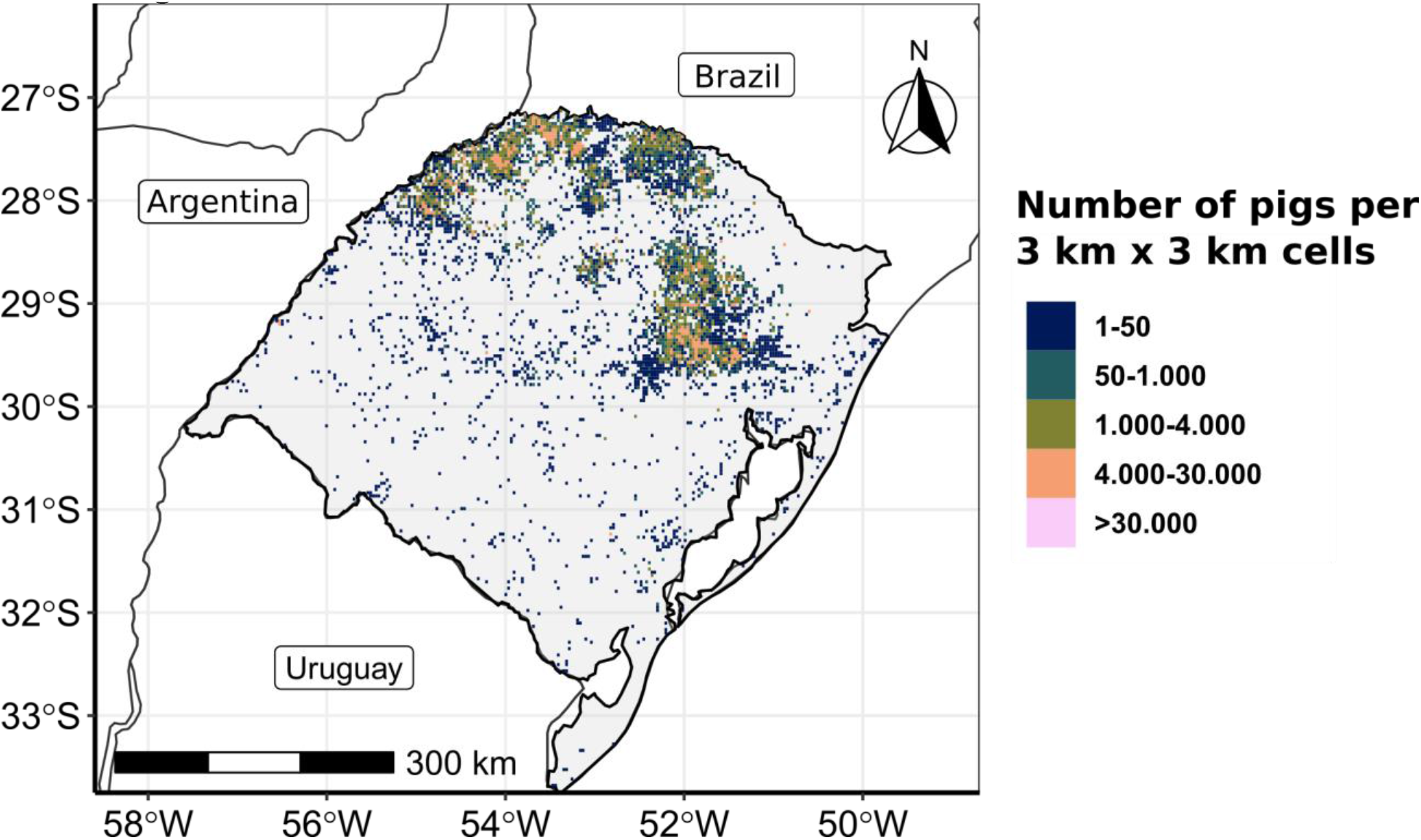
Study area. The study area, Rio Grande do Sul Brazil, was gridded into 3 km x 3 km cells (pixels) to represent the total number of pigs per pixel.

### Modelling ASF transmission dynamics and control strategies

#### Transmission model formulation at the pixel scale

We developed a network model to simulate ASF transmission on a pixel-level directed temporal (daily) network of animal movements. Each pixel (*b* ∈ {1: 4367})can be in one of four health states: susceptible (S), exposed (E), infectious (I), and (Q) quarantined. Details of model parameters are described in Table 1 and a graphical description of disease states and their transitions is shown in Figure 2. When the model initializes, a single pixel is seeded with ASF at a randomly selected daily time point (*t* ∈ *T*) and ends once a pixel is quarantined (see session below). The pixel, the index case, immediately transitions to the infectious state (I). Transmission is then driven by outgoing animal movement between-farms among pixels (between-farm pig movements derived from the SEAPI-RS database (SEAPI-RS, 2018)) from infectious pixels to susceptible ones, and assessed at each sequential time point on a daily scale. We assume that ASF is successfully transmitted from infectious pixels to susceptible ones at a rate ρ (Table 1). When successful transmission occurs, susceptible pixels transition to exposed immediately, and remain in this state for *η*_*b*_days, where *η*_*b*_ ∈ PERT(0, 5.5, 7) and represents the likely incubation period range previously described for high and moderately-virulent ASF strains (Table 1) (Gulenkin et al., 2011; Beltran-Alcrudo et al., 2017). In our model, we assume that the ASF incubation and latent periods are equivalent. The highly virulent strain mimics the dynamics of the current strain circulation in Europe and Asia, Genotype II (Ge et al., 2018; Mighell and Ward, 2021), while the moderate strain is used to represent a subacute form of the disease. We allow *η*_*b*_ to potentially equal zero (i.e., pixels spend zero days in the exposed state and immediately transition to infectious on the following time step) to effectively allow for the possibility that pigs that have already passed their incubation period can be shipped from infected pixels (Howey et al., 2013; Pietschmann et al., 2015). Exposed pixels transition to the infectious state after *η*_*b*_days, and the simulation continues until the first pixel transitions to the quarantined state. The time pixels spent in the infectious state is variable and depends on within-pixel ASF dynamics (Table 1). Quarantine is triggered when a predefined within-pixel pig mortality is reached. Several mortality trigger scenarios were used to start the deployment of control protocols simulated (Table 1 and Figure 2).

**Table 1.** Overview of variables and key model parameters. All units are days unless otherwise noted

| Parameters/<br>Variables | Description | Model values |
| --- | --- | --- |
| T | Vector of sequential timesteps. | 1:1,440 days |
| $\rho$ | The probability that an animal movement from an infected pixel to a susceptible one will cause the latter to transition to the exposed state. | 50%*<br>80%*<br>100% |
| $\eta$ | Vector of likely incubation-period length for acute ASF serotypes (Gulenkin et al., 2011; Beltran-Alcrudo et al., 2017). | 4:7 days |
| $\eta_b$ | Length of ASF incubation period (days), drawn from the beta-PERT distribution (Vose, 2008). The minimum, mode, and maximum values used to parameterize the distribution are taken to be 0, 5.5, and 7. The latter two values represent the mean and maximum of the $\eta$ vector, respectively. | $\eta_b \in \text{PERT}(0, 5.5, 7)$ |
| $\gamma$ | Vector of likely time-to-mortality for acute ASF serotypes (Beltran-Alcrudo et al., 2017; Guinat et al., 2018). Varies with high and moderate virulence. | $\gamma_{high} = 6:9$<br>$\gamma_{mod} = 11:15$ |
| $\gamma_b$ | Time-to-mortality (days), drawn from the beta-PERT distribution (Vose, 2008) parameterized with $\gamma$ values. This is only relevant when the simulation mortality trigger is $> 0\%$. The minimum, mode, and maximum values used to parameterize the distribution are taken to be 0, $\bar{\gamma} = \begin{cases} 7.5 & \text{if } \gamma \in \gamma_{high} \\ 13 & \text{if } \gamma \in \gamma_{mod} \end{cases}$, and $\bar{\gamma}_{max} = \begin{cases} 7.5 & \text{if } \gamma \in \gamma_{high} \\ 13 & \text{if } \gamma \in \gamma_{mod} \end{cases}$ | $\gamma_b \in \text{PERT}(0, \bar{\gamma}, \bar{\gamma}_{max})$ |
| $\beta_{sub}$ | Daily within-pixel ASF transmission coefficient. This is the weighted median of estimated betas presented by (Guinat et al., 2018; Hayes et al., 2020), and first used by (Ferdousi et al., 2019) in their ASF transmission model. This coefficient is only used in the within-pixel model. | 1.679 |
| Virulence | Variable modulating time-to-mortality values. | High, Moderate |
| Mortality trigger | The proportion of the within-pixel population mortality required to trigger quarantine and intervention scenarios. | 0%<br>1%<br>3%<br>10% |
| **Control strategies** | **Description** | **Model values** |
| Quarantine protocol | Procedures for subsequently quarantining pixels nearby or related to those that initially transition from the infectious state.<br>1) **Buffer:** quarantine pixels within a pre-specified distance from an edge of the first quarantined pixel (control zone)<br>2) **Pig production system-wide:** quarantine pixels sharing a production system with the first quarantined.<br>3) **Buffer & system-wide:** a combination of buffer and system-wide procedures.<br>4) **Recent animal movement between pixels:** quarantine pixels with any edges connected to the first quarantined pixel within a pre-specified time period. | 1) buffer<br>2) system-wide<br>3) buffer & system-wide<br>4) recent animal movement between pixels |
| Buffer distance | Distance from initially-quarantined pixels within which other pixels will be additionally quarantined. | 0 km<br>5 km<br>10 km<br>15 km |
| Transfer time frame | Number of time steps (days) within which any observed edges between non-quarantined pixels and initially-quarantined ones will trigger quarantine transition. | 1 day<br>5 days<br>10 days<br>15 days<br>30 days |

\*These $\rho$ values were only used to parameterize simulations intended for sensitivity analyses reported in Supplemental Material and were excluded from all simulations described in the main text.

**Figure 2.**
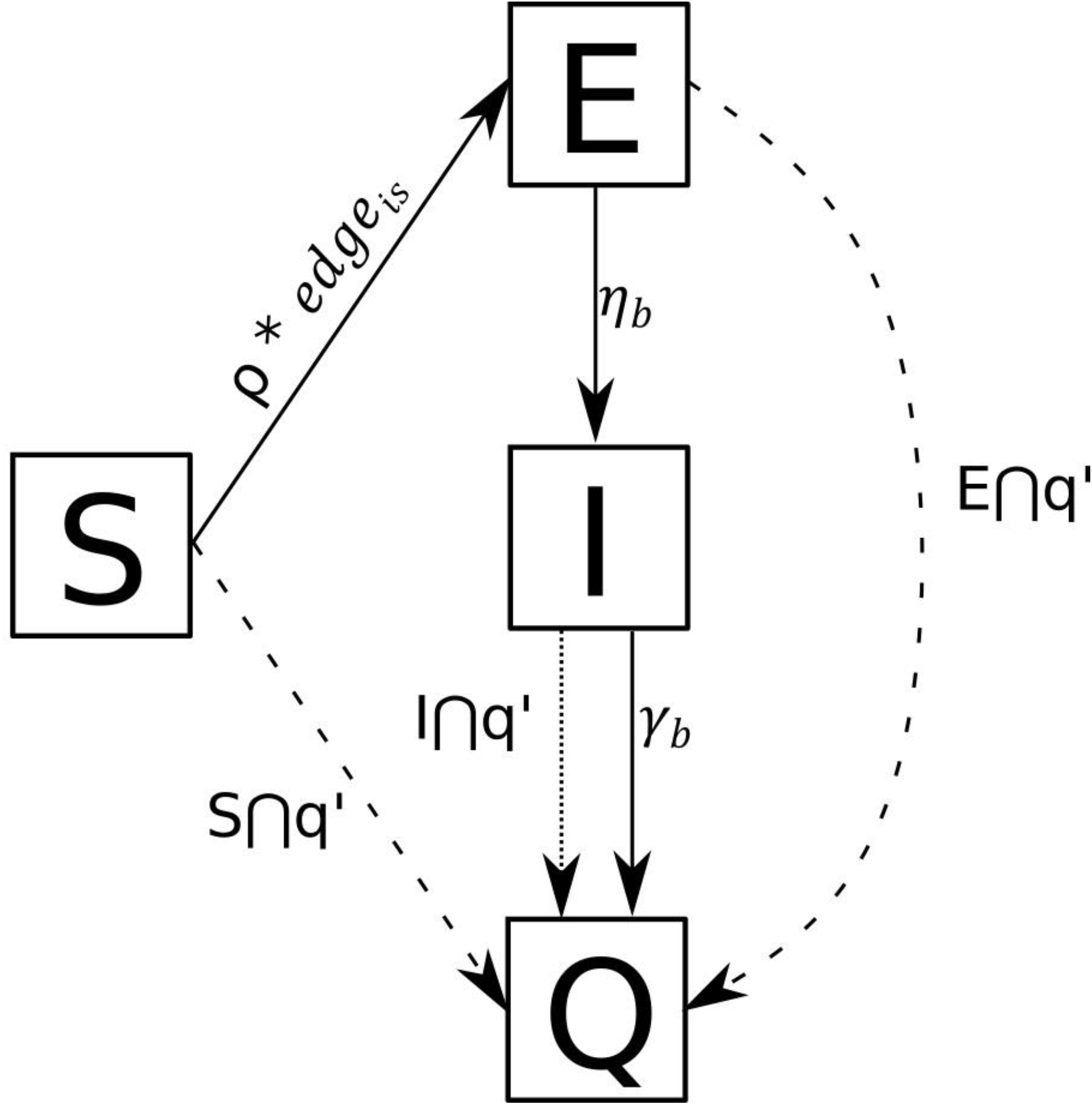
Schematic of state transitions for between pixels model. Solid lines indicate fixed transitions that are bound to occur, dashed lines represent potential transitions that may occur upon quarantine protocols. Pixels in the susceptible state (*s* ∈ S) will transition to the Exposed state with a probability of ρ if an edge exists between susceptible pixel *s* and Infectious pixel *i* (*i* ∈ I) during a defined time period in our dynamic contact network. Pixels in the Exposed state transition to the Infectious state at a fixed rate, and pixels in the Infectious state transition to Quarantined at a fixed rate, as well. Pixels in any state can transition to Quarantined in accordance with a pre-determined quarantine protocol if they share a specific relationship (e.g., are located within a specified radial distance, contain farms operated by the same production system, or have received animals within a predefined time period) with the first pixel to transition from I to Q (i.e., q′).

When mortality trigger was set to be zero (i.e., nearly perfect disease detection), named hereafter as “optimistic scenario,” in which we assumed that infectious pixels transition to quarantined as soon as the first animal dies, which occurs after *γ*_*b*_days, where *γ*_*b*_ ∈ PERT(0, 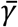, max(γ)). The γ parameter is a vector describing the likely time-to-mortality range based on (Beltran-Alcrudo et al., 2017; Guinat et al., 2018), and like *η*_*b*_can equal zero to simulate instances when ASF leads to severe clinical infections assumed to be more rapidly detected by passive or active surveillance. When the mortality trigger was above zero, we used a within-pixel compartmental Susceptible-Exposed-Infectious-Recovered (SEIR) sub-model describing ASF transmission in local swine populations to estimate the number of days required for mortality trigger percentage of animals within a pixel to die from ASF, and set *γ*_*b*_ to this value. The sub-model is given in equations:

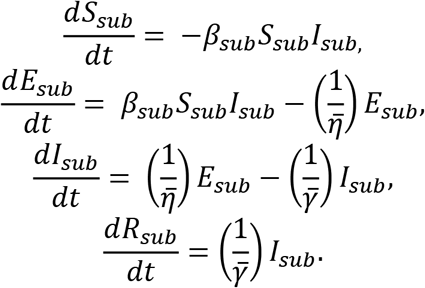

Note that the “sub” designation denotes components of the sub-model, and *R*_*sub*_ here can be considered to be dead individuals. We assumed that the number of pigs (*N*) that a pixel contained was equal to the maximum swine-holding capacity of all farms within a pixel, and this sub-model was always initialized with a single infectious pig and *N–1* susceptible pigs. When mortality trigger is above zero, infectious pixels transition after the minimum number of days required for mortality trigger percentage of within-pixel swine populations to be removed from the *I*_*sub*_compartment 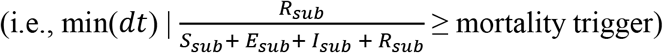. When infectious pixels transition to Quarantined, additional pixels may be similarly transitioned, regardless of their current state at time step *t*, in accordance with spatially-explicit quarantine protocols (Table 1). For example, control zones of varying sizes (e.g., 1-km, 10-km, or 15-km radii) can be simulated to quarantine all pixels within a fixed distance from the infected pixel that initially transitioned. All pixels quarantined due to their relationship (e.g., pixels with at least one farm of the same pig producing company), pig shipping history, or distance from other pixels were recorded as such in simulation output to support the evaluation of quarantine effects on nearby areas.

We make a number of assumptions in our sub-model. First, we assume that the modeled ASF serotype induces acute symptoms in infected pigs (i.e., our parameterization is not based on per-acute or chronic signs), and infections will always cause death. Second, we assume that local swine populations are static within pixels, regardless of any observed animal movements between-farms. If no information is known about an infectious pixel’s swine population or it was reported as zero, the rate at which it transitions to Quarantined is determined as if the mortality trigger is equal to zero, regardless of the parameter value of the simulation. For example, even if the mortality trigger parameter for a simulation is set to 3% of pixels’ swine populations, if no information is known about this population for any given infected pixel, the time-to-quarantine following infection (*γ*_*b*_) will simply be drawn from the previously-described beta-PERT distribution. Finally, we assume no external transmission sources (e.g., spillover from feral swine populations) contribute to ASF exposure.

### The effect of cumulative mortality triggers and control strategies scenarios

Four delayed ASF control triggers based on mortality were simulated individually for each ASF strain (highly and moderately virulent), and later combined with quarantine base control scenarios (Table 1). In the model, ASF infection was detected based on cumulative mortality assumed to be identified by passive surveillance, the cumulative proportion of dead pigs within the simulated thresholds, or in combination with active surveillance from veterinarians or official services investigations on farms with clinical signs or cumulative mortality that trigger an epidemiological investigation, described in Table 1. Model simulations were run across a factorial combination of every variable combination, ASF strains, four mortality triggers, 0%; 1%; 3%; 10%, and the following control measures: 1) established control zones, buffers of a minimum from 0 km to 15 km radius surrounding the affected pixels; 2) implementation of quarantine to all farms of a pig producing company, named hereafter as system-wise, linked to pixels either directly quarantine or infected; 3) backward tracing pixels with direct pig movements to or from pixels under quarantine or infected; and 4) the combination of control zones and the system-wise quarantine (see Supplementary Figure S2 for a graphic representation control scenarios). Model simulations factorial combination were the following:

1. High and moderate virulent ASF strains & four mortality triggers (0%; 1%; 3%; 10%) & one quarantine protocol (system-wide).
2. High and moderate virulent ASF strains & four mortality triggers (0%; 1%; 3%; 10%) & 4 buffer distances (0 km; 5 km; 10 km; 15 km) & 2 quarantine protocols (buffer; buffer & system-wide).
3. High and moderate virulent ASF strains & four mortality triggers (0%; 1%; 3%; 10%) & five between pixel animal transfer timeframe levels (1 day; 5 days; 10 days; 15 days; 30 days) & 1 quarantine protocol (recent between-pixel animal movements).

### Model outputs and analyses

Each simulation produced three outputs: i) pixel state variables: Susceptible, Exposed, Infectious, and Quarantined, at the final time step; ii) maximum transmission distance and epidemic duration; and iii) the observed transmission and quarantine networks (i.e., who infected or triggered quarantine for whom, and at what time step). As measures of potential severity, we calculated the duration in days, spread capacity in kilometers, and secondary attack rates of ASF epidemics prior to enacted quarantines at the pixel level.

Furthermore, for each quarantine protocol, we report and compare the probability that enacted quarantines resulted in complete ASF containment, (i.e., all pixels observed in the Infectious or Exposed state at any point in the simulation transitioned to the Quarantined state at the final timestep), and the mean number of susceptible and “infected” (i.e., exposed of infectious) pixels quarantined. In addition, we derived the total number of quarantined uninfected (i.e., unexposed entities that were quarantined due to their relationship with, or proximity to, a quarantined infected pixel) and infected pixels, as well as the expected number of farms and pigs affected according to each factorial combination. In a series of choropleth maps, we summarize pixel-level probabilities of being infected as a number of secondary cases.

The simulation procedure used each 3-km pixel containing ≥ 1 farm (n = 4,367) in the animal shipment network as the initially-seeded infectious pixel 50 times, resulting in 24,455,200 unique simulations. This number of iterations was deemed sufficient to obtain stable outcomes, based on calculations for estimating the appropriate number of simulations in Monte Carlo analyses (Supplementary Material Table S1). In addition, a sensitivity analysis was performed by varying the probability that animal movements from an infected pixel to a susceptible one resulted in a successful infection event, ρ (Table 1 and Supplementary Material) and results are available in the Supplementary Materials. The model was developed in the R (3.6.0 R Core Team, Vienna, Austria) environment, and all simulations were run in RStudio Pro (1.2.5033, RStudio Team, Boston, MA). More details about model implementation can be found in this public repository (https://github.com/machado-lab/ASF_model)

## Results

Approximately 87% of simulations resulted in no secondary infections because no animal movement edges from the initially-seeded pixel existed in the daily pixel-level networks from the start of simulations to the quarantine time of the initial pixel. Thus, in these cases ASF could not propagate between pixels. The epidemic duration and number of secondary cases were positively correlated with the mortality trigger, while the maximum distance spread was not determined by mortality triggers (Figure 3). The number of days to contain ASF under the optimistic control scenario (0% mortality trigger) for the moderately and the highly virulent strain was, on average 11.2 (sd: 2.49) days and 6.50 (sd: 1.59) days, respectively. At 10% mortality the epidemic duration mean was 15.9 (sd: 5.55) days and 13.1 (sd: 7.22) days, respectively (Figure 3; Supplementary Table S2). The average number of secondary cases in simulations was 1.53 (sd: 2.08) and 1.25 (sd: 1.03) pixels at the optimistic scenario (0% mortality trigger), and at 10% mortality 2.75 (sd: 6.71) and 2.56 (sd: 5.95) pixels for the moderate and the highly virulent strain, respectively (Figure 3; Supplementary Table S3). Finally, we also found that the spread distance from initially-infected pixels was similar between both ASF strain variants, on average 23.8 km (sd: 68.4) and 17.9 km (sd: 59.7) in the optimistic scenario (0% mortality trigger) and at 10% mortality disease control trigger ASF spread could stretch up to 231.1 km (sd: 78.2) and 29.5 km (sd: 76.0), for the moderately and the highly virulent strain, respectively (Figure 3; Supplementary Table S4).

**Figure 3.**
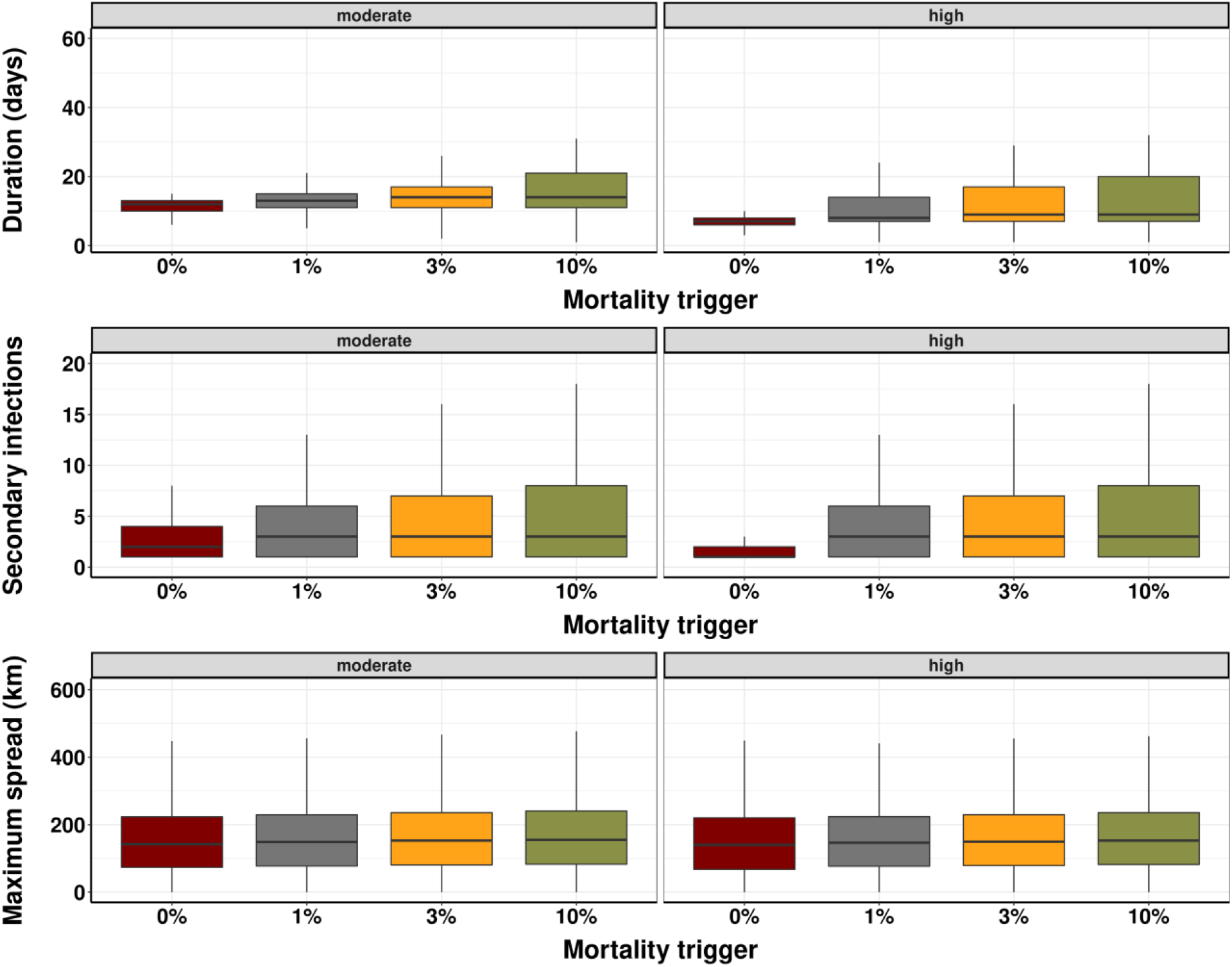
ASF epidemic outcomes for a highly or moderately virulent strain. The predicted number of days in an ASF epidemic using different mortality levels to trigger control interventions (top). The predicted number of secondarily infected pixels (middle). The maximum distance from initial seeded infection to all secondary infections during the course of the simulations (bottom). The boxplot displays in the vertical line the maximum value, while horizontal line the median and the top limit of the box the third quartile and the bottom box limit the first quartile.

The predicted spatial distribution pattern of an ASF epidemic is illustrated in Figure 4. An excess number of secondary cases were estimated to occur in densely populated locations in the north and northeast regions of the state. There was no evidence of significant variation in the spatial distribution of ASF strain-specific distribution (Figure 4). However, as expected, early response resulted in fewer successful propagation, especially when the control triggers were activated as soon as the first ASF was found (Figure 4, Supplementary Figure S3 for 3% and 5% mortality trigger maps).

**Figure 4.**
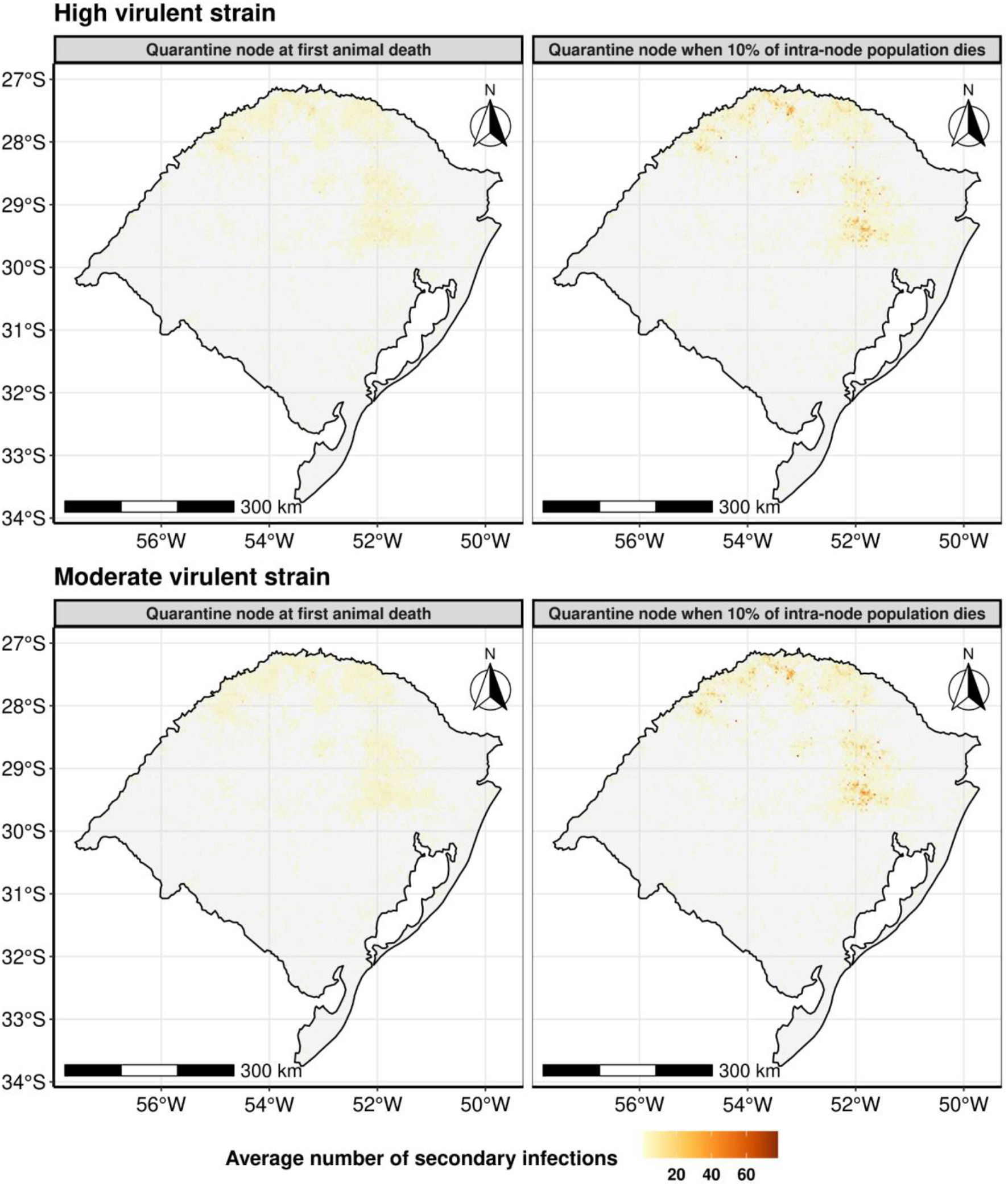
Average infected pixels (3 km by 3 km). Pixels highlighted correspond to simulated average secondary cases under the simulated optimistic scenario (quarantine is triggered as the first ASF case is identified), left panel, compared with a delayed response trigger after a cumulative 10% of the pig mortality, right panel.

Under the model considered, if control actions are triggered t as the first animal dies, we predicted ASF epidemics will lead to an average of 18.79 (sd: 120) farms and 2,246 (sd: 7,525) pigs for a highly virulent strain and 23.77 (sd: 137) farms and 3,466 (sd: 12,318) pigs for a moderately virulent strain. In comparison, with a mortality based delay of 10% of the population, the epidemic was expected to be nearly double the size, with 42.97 (sd: 120) farms and 7,927 (sd: 7,524) pigs for a highly virulent strain and 46.28 (sd: 137) farms and 8,669 (sd: 12,318) pigs for a moderately virulent strain (Table 2).

**Table 2.** Predicted quarantine consequences of two ASF strain outbreaks under different mortality triggers. The average and standard deviation in brackets of the predicted quarantined pixels, farms, and animals are shown.

| ASF strain virulence | 0% Mortality trigger |  |  | 1% Mortality trigger |  |  | 3% Mortality trigger |  |  | 10% Mortality trigger |  |  |
| --- | --- | --- | --- | --- | --- | --- | --- | --- | --- | --- | --- | --- |
|  | *Pix els* | *Farm* | *Pigs* | *Pix els* | *Far m* | *Pigs* | *Pix els* | *Far m* | *Pigs* | *Pix els* | *Far m* | *Pigs* |
| Mode rate | 1.53<br>(2.08) | 23.77<br>(136.75) | 3,466.37<br>(12,318) | 2.01<br>(4.1) | 32.62<br>(170.48) | 5,573.08<br>(21,744.84) | 2.35<br>(5.13) | 37.9<br>(189.47) | 6,789.7<br>(26,226.59) | 2.75<br>(2.08) | 46.27<br>(136.74) | 8,668.13<br>(12,317.84) |
| High | 1.25<br>(1.03) | 18.79<br>(119.7) | 2,246.59<br>(7,525.49) | 1.92<br>(3.77) | 31.06<br>(164.64) | 5,220.32<br>(20,408.78) | 2.19<br>(4.69) | 35.9<br>(181.94) | 6,353.88<br>(24,453.79) | 2.56<br>(1.03) | 42.96<br>(119.7) | 7,926.09<br>(7,523.99) |

### Mortality-based response triggers and effectiveness of control strategies

Our model also allowed for the investigation of complete disease control (Figure 5; Supplementary Table S5). The less effective strategy across all scenarios was a buffer of less than 15 km. The 15 km buffer outperformed the system-wide quarantine, and ≥ 10 km buffers by themselves outperformed the 1-day shipment trace back. Additionally, when the buffers were used in conjunction with the system-wide control, efficacy increased. The rank-order of control strategies were the same for the both virulent strain and mortality-triggers scenarios (Figure 5) Under the optimistic mortality-base response trigger (0% mortality trigger), 95.5-to-99.5% of simulations wherein pixels that had contact with the initially-quarantined pixel in the previous 15-to-30 days were also quarantined resulted in complete containment of the ASF epidemic (Figure 5). When we simulated 10% of pig mortality, 30-day contact tracing resulted in 89.3% and 88.9% containment probabilities, for the high and the moderate virulence strains, respectively (Figure 5 and Supplementary Table S5).

**Figure 5.**
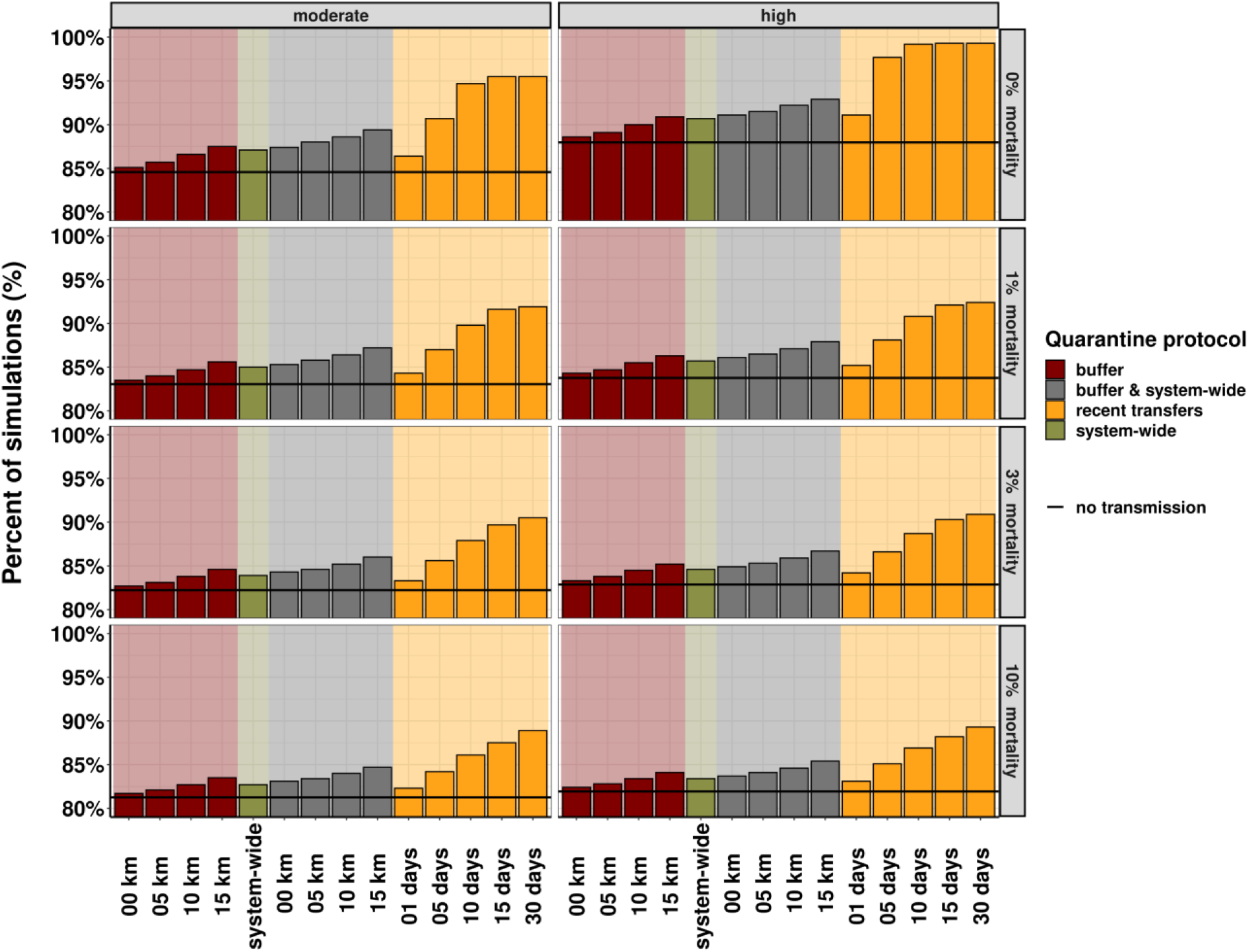
The percent of ASF epidemics successfully halted by mortality-trigger and virulence combinations. The “no transmission” line indicates the percent of simulations in which no secondary transmission occurred because no animal movements were observed from the seeded pixels between simulation start times and the time of quarantine for these cases. Thus, quarantining only the initially-seeded pixel was sufficient to completely contain the outbreak.

The implemented backward contact trace quarantine strategy targeted at infected premises had the lowest total quarantined farms compared with other strategies, regardless of ASF strains and mortality triggers (Figure 6). The overall number of quarantined farms under the 0 km buffer strategy, when compared with 1-day shipment trace back, was similar, the latter control strategy was by far more effective in halting transmission. The number of infected farms quarantined under the 0 km buffer was on average 15.07 farms, similar to one day of recent backward contact trace 16.06 farms (Supplementary Table S6). Whilst the number of susceptible (uninfected) farms quarantined was on average 79.22 farms for the 0 km buffer, while under 1-day shipment trace back exhibited on average 38.10 farms. Overall, the least negative economic and animal welfare impact quarantine strategy was to utilize backwards contact tracing-based. For example the 30-days scenario and at 10% mortality-trigger with highly virulent strain, only 16 of uninfected farms were predicted to be quarantined, while about 1,258 of the total uninfected farms would be quarantined via a system-wide control strategy, for example (Supplementary Table S6). For the full details of the simulated quarantine, strategies, including the success of each intervention on complete epidemic containment, see Supplementary Table S6, which also includes results for pixels and the number of pigs predicted to become quarantined.

**Figure 6.**
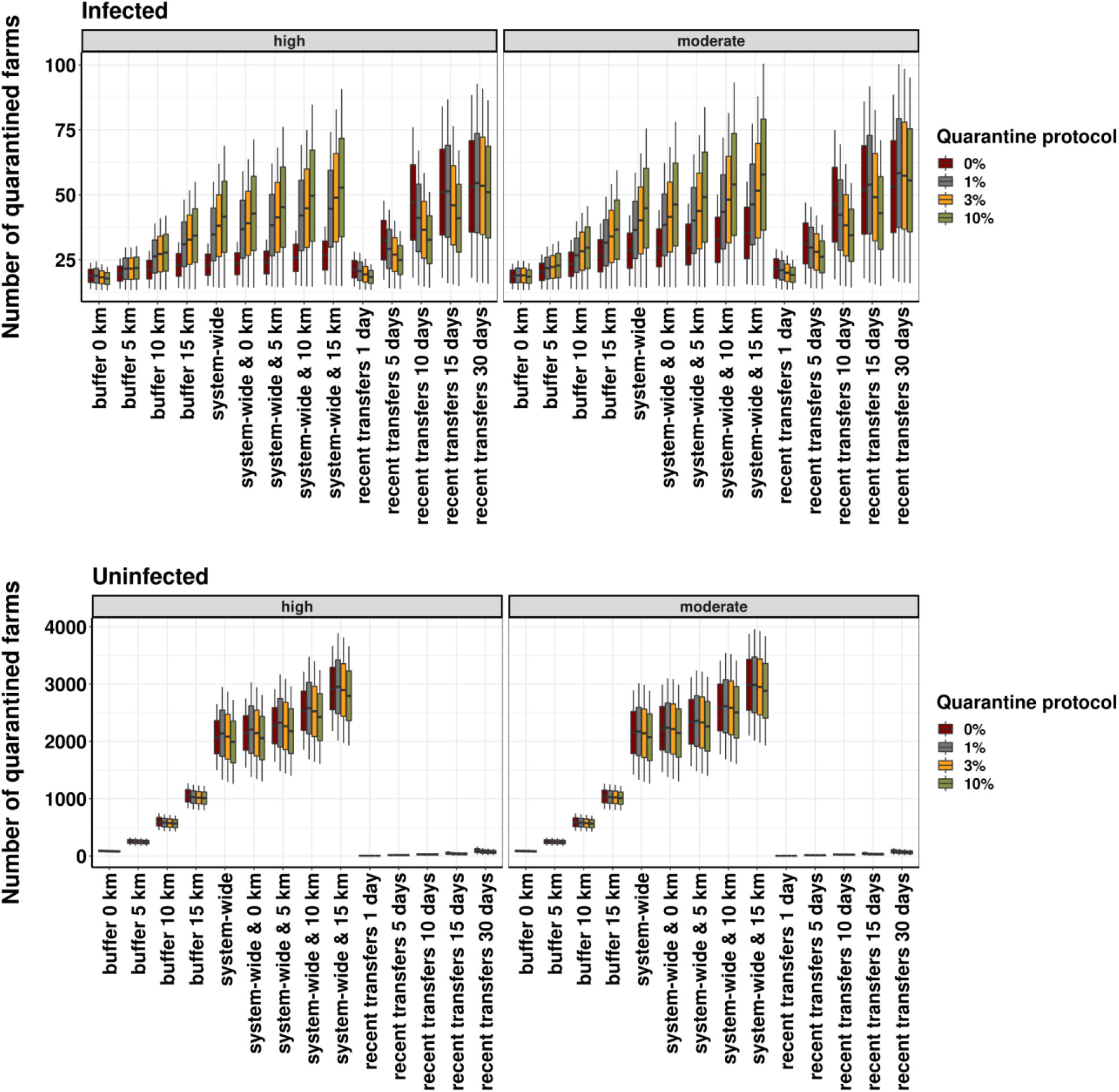
The predicted number of quarantined farms per mortality-trigger, ASF strain, control strategies, and infected and uninfected farms. Summary of simulated outputs the total number of quarantined farms, regardless of its infection state. Table S6 included the breakdown of infected and uninfected states of the quarantined pixels, farms, and number of pigs.

## Discussion

In ASF-free countries, the lack of experience containing large-scale disease emergencies limits the development of evidence-based disease control programs and more precise contingency plans (Halasa et al., 2016b, 2018a; Brown et al., 2020). In this context, we developed a nested multiscale model to simulate the spread of moderately and highly virulent ASF strains in Brazil (Beltran-Alcrudo et al., 2017; Guinat et al., 2018). We demonstrated that epidemic duration as expected was prolonged and the number of secondary infections was higher under the introduction of a moderately virulent strain. Our model predicted that delaying disease control response under a 10% mortality trigger resulted in an epidemic duration and number of secondary cases that were more than double that of the “optimistic scenario” in which disease control started on the same day that the first dead pig was identified. The most effective simulated intervention strategy, tracing and quarantining previous contacts for the previous 15 or 30 days had nearly the impact in reducing ASF propagation. Our analysis on the burden caused by the implemented quarantine strategies points again to the advantages of backward contact tracing in which the minimum number of farms were predicted to be quarantined. In addition, our model showed significantly fewer uninfected (not infected states in the model simulation) pixels were quarantined under backward contact tracing protocol than after implementing any other intervention (Supplementary Table S6).

The simulated epidemic durations were relatively short and directly correlated with the delayed mortality-based control triggers (Figure 3). Our predicted epidemic duration peaked at a maximum of 15 days, similar to other simulated work in which the median epidemic duration was 20 days (min 6-47 days) (Andraud et al., 2019), and with the Denmark work that estimated it to be between 1 to 29 days (Halasa et al., 2016a). The vast majority of the simulations did not result in large outbreaks, the average number of secondary cases varied from 18.79 farms to 46.27 farms depending upon the simulated ASF strain, with limited long-distance spread, which has also been observed elsewhere (Andraud et al., 2019; Akhmetzhanov et al., 2020). One reason for the similarity could be associated with pig population structure and much less vertically integrated pig farming in the EU and in Brazil when compared with the highly integrated production systems in North America (Galvis, Prada et al., 2021). Another commonality with the simulation done in France was the distances from seed infection to infected pixels estimated to be 300 km (Andraud et al., 2019), while in our study it was restricted to an average distance of 30 km with a maximum spread distance of 603 km (Table S4). In support of our results, China 2018-2019 ASF outbreaks also reported. Furthermore, the long distance spread described in our study was likely related to the movement of pig among independent pig producers. Different from farms producing pigs to commercial pig companies which only contract with site within a predefine distance to slaughterhouses and or feed mills, independent or mid- and small- scale producers have more freedom sell or transport animals to much greater distances. Indeed, movement among stallholders has been demonstrated in other countries such as in China, with documented ASF transmission by the transportation of animal between farms located more from 200 km to 500 km apart (Akhmetzhanov et al., 2020). Future studies are needed to explore such epidemic dynamics in more vertical integrated systems such as in the U.S. (Galvis et al., 2020), to better evaluate the role of the effects of heavily integrated pig production (i.e., North America) in the dissemination of ASF while also evaluating the feasibility of control and eradication counter measures (Halasa, Ward et al., 2020).

We provided the first mapping of ASF epidemic potential in Brazil showing the number of predicted secondary infections, which highlighted areas of a high density of commercial sites with the greatest potential of spread. Similar spatially explicit disease spread models have been developed to study the spatial distribution of the probability of epidemics for Classical Swine Fever in Great Britain (Porphyre et al., 2017), and also in Bulgaria (Martínez-López et al., 2013), both also concluded that distinct high-risk areas were concentrated in pig-dense regions. While the development of ASF control programs and strategies remains a work in progress for affected countries (European Food Safety Authority (EFSA) et al., 2018; Miller and Pepin, 2019), in practice, the maps produced here could be used directly by the local swine industry and by the official veterinary services to guide surveillance activities. An added value of mapping could include, for example, the development of wildlife risk based passive surveillance of wild boar carcass identification and removal, while for domestic pig production such high risk areas could be used as target for syndromic surveillance which could include pig mortality as the main trigger.

The results suggested that 30 days backward contact tracing and deployment of quarantine to all movements from pixels in contact with infected or exposed pixels, was the strategy of choice. We could argue that one of the reasons for a better performance of backward contact tracing could be associated with the model mechanics in which explicit local spread is not formally considered, besides the consideration of the sub-model at pixel level. Because modelling results suggest that if control tactics are activated at this first case is detected there is nearly not advantages to contact trace for more than 5 days. However, delay in detection is expected, therefore the implementation of backward contact trace increase clearly show the reduction in secondary cases as mortality trigger increased (Figure 5). Similarly to our model, a work done in France, in which the structure of pig production with many independent pig producers has also concluded that 99% of ASF transmission resulted from between-farm pig movement, therefore centering control strategies at this mode of transmission compared with quarantine control zones is more likely to reduce ASF propagation (Andraud et al., 2019). Even though we have not directly modeled local transmission (farm-to-farm proximity), our sub-model considers ASF spread dynamics within a pixel’s total pig population and implicitly captures unmeasured processes such as shared equipment, tools, and people between neighboring farms within pixels (Halasa et al., 2016a). In addition, our model results reiterate the recommendation of (Halasa et al., 2018a), in which increasing control zones are not expected to significantly shorten ASF epidemics. Additionally, deploying control zones even at 5 km may not be feasible in areas with high pig population densities, such as the northeast of Rio Grande do Sul, which encompasses an average of 15 farms within 3 km x 3 km cells (pixels). Likewise, permitting animal movement during and after an ASF outbreak is detected becomes an extremely large and complex task to manage, thus indeed add another burdern to the official veterinary services, on top of the laboratory capacity needed for non-infected sites requesting movement permits. Finally, we demonstrated multiple control methods when applied in parallel (i.e., control zone and backwards contact trace) had less than 5% improvement in the control of the ASF propagation.

Pre-emptive depopulation has been described as an ASF control measure in a number of simulation studies and performed during ASF outbreaks, and has proven to be considerably effective in flattening the curve of ASF epidemics (Halasa et al., 2018b; Faverjon et al., 2020). However, pre-emptive depopulation of healthy animals would be met with considerable resistance from the general public and pig producers. Our model predicts that 30-day contact trace would unnecessarily quarantine 16 uninfected farms, while 15 km ring-based quarantine was predicted to massively increase the direct cost by the inflated number of farms needed to undergo quarantine was estimated to be 797 farms, see Table S6 for the results of all simulated strategies. Given the real time and electronic movement recorders implemented in the region, contact trace would be the preferred first control strategy in the ASF control program. Even though immediate culling of all pigs at sites where ASF is detected, as well as blanket depopulation within 3-km epidemic zones around infected pig sites (e.g., farms, backyards), used by Chinese authorities, (FAO, 2019c), blanket depopulation based on geospatial proximity is unlikely to stop transmission (Akhmetzhanov et al., 2020).

### Limitations and further remarks

We identify a number of limitations in our study associated with simulation rule decisions and data availability. First, our model unit (3-km^2^ pixels) targeted spatial locations where infection would spread and move quickly, thus it was not our objective to make inferences at the farm level. Ultimately, model outputs match the local authorities surveillance system which has been directed towards high and low disease transmission areas. In addition, it was not our goal to estimate the contribution of the route of transmission, which could include local transmission, airborne on a smaller scale (de Carvalho Ferreira et al., 2013; Galvis, Jones et al., 2021). However, the next steps could include other routes of transmission including the indirect contact of feed delivery and potential reinfection. Given the importance of wild boars in the spread of ASF, we recognize that there is a need to further develop simulation models to also include the involvement of ASF-outbreaks in wild boar (Hayes et al., 2020).

We simulate ASF spread via pig movements between premises registered with the state veterinary services, which is undoubtedly associated with a large proportion of ASF outbreaks (Andraud et al., 2019; Gao et al., 2020). The potential effects of animal movements from small “backyard” farms on ASF transmission are discounted in our model. It is plausible that the current local surveillance capacity is not sensitive enough to detect small mortality proportions, particularly in small pig farms, future studies considering not only backyard pig movement data but also the sensitivity of disease surveillance should be considered. Similarly, because our data only include farm locations and pig movements within the state of Rio Grande do Sul, potential effects of multi-state animal shipments on ASF spread are also masked from our analyses. Future studies should also strive to make shipment networks as complete as possible.

Another limitation that must be addressed regards our decision to end simulations as soon as any agent transitions to the quarantined state. In reality, pigs would still continue to be moved throughout the network following the quarantine of one or many areas, and within-pixel transmission may continue to occur even after quarantine protocols are deployed. Furthermore, we would expect a multi-day response time when quarantining locations, regardless of quarantine protocol (e.g., distance-based quarantine). In our model, however, we do not make any assumptions or predictions regarding how our animal shipment network will behave once pixels are quarantined. Thus, we do not simulate ASF transmission post-quarantine, and we do not allow multi-day response times for triggering quarantine for pixels associated with the initially-quarantined one. Another limitation to be noted is how the mortality-trigger was simulated to all related with ASF mortality, this over simplification was assumed because we lack data of mortality caused by other infectious diseases or production related events. Therefore, more information about the sensitivity of current mortality detection systems is needed to reduce uncertainty about the feasibility and effectiveness of the modelled mortality rate. Finally, it was beyond our aims to consider economic impacts of epidemics, therefore future work could consider the full evaluation of direct and indirect costs of losses with mortality and stepping out, while also considering the burden of the official veterinary services including diagnostic and movement permits management.

Our model demonstrates which areas are at consistently high risk for pig-infections. The results could be used to predefine risk areas prior to the introduction of ASF but also more broadly other foreign animal diseases (FADs) (Gao et al., 2020). In addition, it is pivotal that not only the official veterinary services are aware of the risk distribution of ASF but also that the pig industry receives appropriate information in order to jointly prepare. Given the absence of effective vaccines, the only available measure relies on improving on-farm biosecurity. Thus far there is no clear regulation for the control of ASF-outbreaks in Brazil, but contingency plans against FADs demand mass depopulation of infected farms and the investigation of all herds within the surveillance zones. Even though recent outbreaks in Europe, Estonia, showed that for large pig farms mortality is not the best option for early detections of an ASF outbreak (Nurmoja et al., 2020). Future studies are needed, for example considering the mortality levels at barn and pen-level (Faverjon et al., 2020). It is worth noting that even though fatality rates under field conditions have been high, while initial mortality, depending on the ASF strain, have been reported to be low in especially in large commercial sites (C. et al., 2018; Lamberga et al., 2020; Nurmoja et al., 2020), under such scenarios it could be mistaken for other endemic diseases such as porcine reproductive and respiratory syndrome (Faverjon et al., 2020).

## Conclusion

The vast majority of simulations predicted short epidemic durations and resulted in relatively small outbreaks. Early mortality-based ASF detection significantly curbed the epidemic size, especially when a best-case scenario, in which quarantine is applied as the first infected pig dies, is compared with 10% pig mortality as a quarantine trigger. Our modeling analyses have given insight into transmission dynamics and ASF spatial distribution in which it may persist, thus the probability map of secondary cases could be used to define priority and represent the first steps for well-programmed surveillance. Although our model only considered quarantine as control strategies, the best intervention regardless of delayed disease control triggers was backward contact tracing, in which both 15 and 30 days showed nearly the same efficacy in reducing propagation, importantly even applying quarantine to less contacts (e.g., 5 and 10 days) over performed other interventions. Additional quarantine efforts based either on ring-based strategies or pig production level did not significantly improve control efficacy. Moreover, the best control strategy was independent of viral strain virulence through all simulated scenarios. Finally, the results of this study can be used to make informed decisions at the state of Rio Grande do Sul Brazil to implement target ASF surveillance activities and used in preparation for future outbreaks. Our mapping results showed areas in which ASF spread is more likely, therefore further preparation would be beneficial in such areas.

## Acknowledgments

The primary funding support of this project is from Global Health, College of Veterinary Medicine at North Carolina State University and by the Fundo de Desenvolvimento e Defesa Sanitária Animal (FUNDESA‐RS).

## Authors’ contributions

TF, CL, and GM conceived the study. GM designed the study and coordinated data collection with the assistant of FPNL. TF and GM conducted data processing, cleaning, designed the model, and simulated scenarios with the assistance of CL. TF, CL, and GM designed the computational analysis. TF and GM wrote and edited the manuscript. All authors discussed the results and critically reviewed the manuscript. CL and GM secured the funding.

## Conflict of interest

All authors confirm that there are no conflicts of interest to declare.

## Ethical statement

The authors confirm the ethical policies of the journal, as noted on the journal’s author guidelines page. Since this work did not involve animal sampling nor questionnaire data collection by the researchers there was no need for ethics permits.

## Data Availability Statement

The data that support the findings of this study are not publicly available and are protected by confidential agreements, therefore, are not available.

## Notes

### Competing Interest Statement

The authors have declared no competing interest.

