## supplementary for "Modeling the role of mortality-based response triggers on the effectiveness of African swine fever control strategies"

**Table S1.** The estimated number of simulations required to capture significant differences in model metrics, as reported by three equations. All estimates are rounded up to the nearest whole number.

| Variable | Mean (sd) | Alpha level | Simulation estimate (1)† | Simulation Estimate (2)§ | Pilot |
|---|---|---|---|---|---|
| Epidemic duration (days) | 11.86 (5.31) | 0.05 | 43397 | 481 | 22708320 |
| Maximum epidemic distance (km) | 26.43 (72.18) | 0.05 | $8.00e^{12}$ | $8.87e^{10}$ | 22708320 |
| Number of nodes infected | 2.06 (4.57) | 0.05 | 32061 | 356 | 22708320 |
| The quarantined proportion of infected nodes | 0.92 (0.22) | 0.05 | 75 | 1 | 22708320 |
| Number of healthy nodes quarantined | 36.91 (80.31) | 0.05 | 9910540 | 109813 | 22708320 |

†Estimate 1 was obtained from the equation given by (Winston, 2020). §Estimate 2 was obtained from the equation given by (Díaz-Emparanza, 2002). (Of note, 90 of these simulations extreme values drawn from beta-PERT distributions resulted in epidemic periods > 60 days, which is unrealistic for ASF outbreaks in our study system. These 90 simulations were removed from the data set prior to any analyses. All analyses were therefore carried out on a data set containing 24,455,110 simulations.

**Table S2.** The distribution of the number of days after ASF introduction according to the simulated response speed based on pig mortality. The table presents the mean and standard deviation.

| Time to response (mortality-base) | ASF strains |
|---|---|

| | High | Moderate |
|---|---|---|
| **At the first dead pig** | 6.50 days (sd: 1.56) | 11.2 days (sd: 2.49) |
| **1% mortality** | 10.0 days (sd: 4.40) | 12.7 days (sd: 3.25) |
| **3% mortality** | 11.4 days (sd: 5.63) | 14.1 days (sd: 4.11) |
| **10% mortality** | 13.1 days (sd: 7.22) | 15.9 days (sd: 5.55) |

**Table S3.** The distribution of secondary cases according to the response speed based on pig mortality. The table presents the mean and standard deviation.

| **Time to response (mortality-base)** | **ASF strains** | |
|---|---|---|
| | **High** | **Moderate** |
| **At the first dead pig** | 1.25 (sd: 1.03) | 1.53 (sd: 2.08) |
| **1% mortality** | 1.92 (sd: 3.77) | 2.01 (sd: 4.10) |
| **3% mortality** | 2.19 (sd: 4.691) | 2.30 (sd: 5.13) |
| **10% mortality** | 2.56 (sd: 5.95) | 2.75 (sd: 6.71) |

**Table S4.** The distribution of distance spread as ASF introduction according to the simulated response speed based on pig mortality. The table presents the mean and standard deviation.

| **Time to response (mortality-base)** | **ASF strains** | |
|---|---|---|
| | **High** | **Moderate** |
| **At the first dead pig** | 17.9 km (sd: 59.7) | 23.8 km (sd: 68.4) |
| **1% mortality** | 25.6 km (sd: 70.9) | 27.1 km (sd: 73.0) |
| **3% mortality** | 27.5 km (sd: 73.4) | 28.9 km (sd: 75.4) |
| **10% mortality** | 29.5 km (sd: 76.0) | 31.1 km (sd: 78.2) |

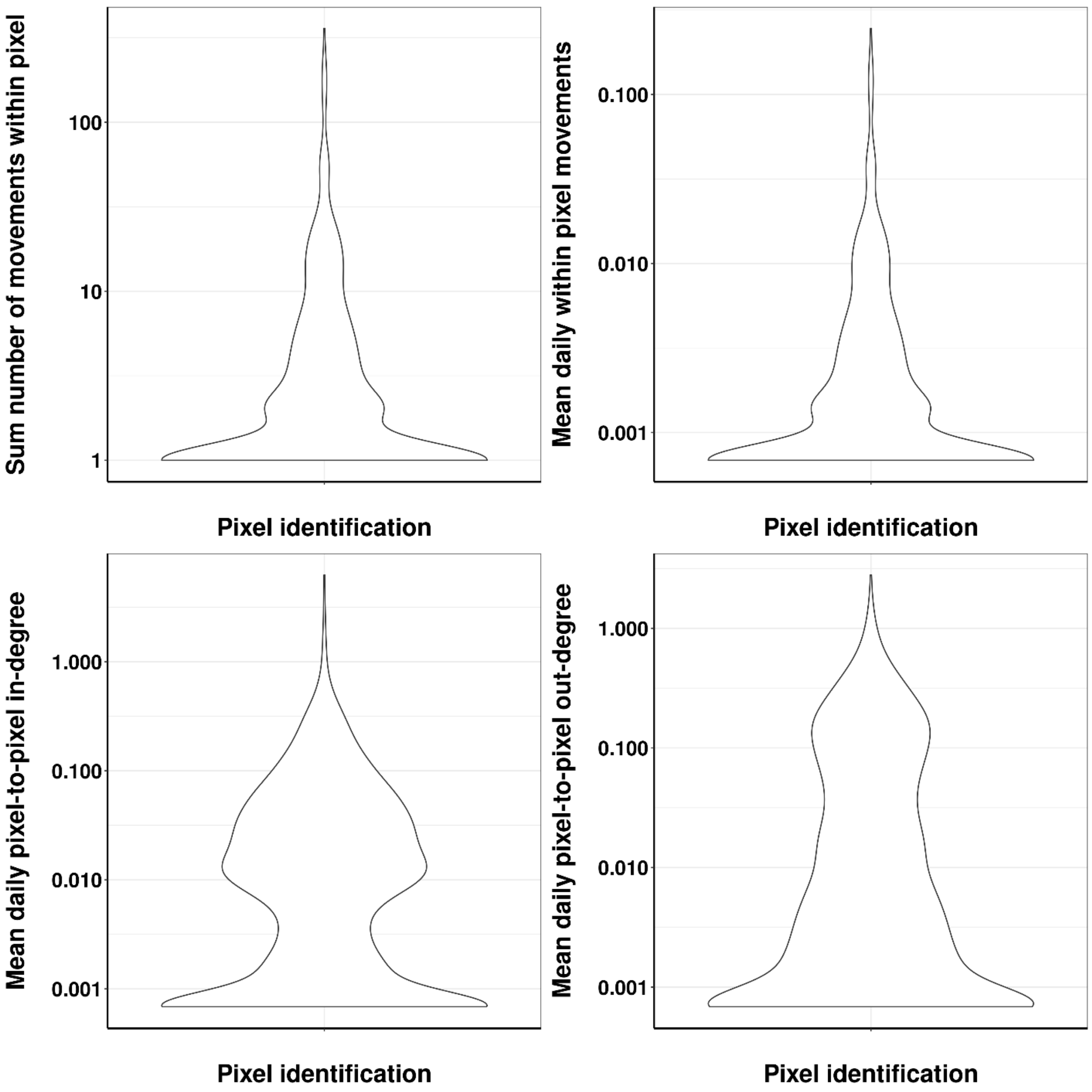


**Figure S1.** In the top panel shows violin distributions of the daily number of movements within the 4,367 unique 3-km pixels (top-panel), while in the bottom-panel violin distribution of the between-pixel movements represented by an in- and out-degree. Y-axis is log 10.

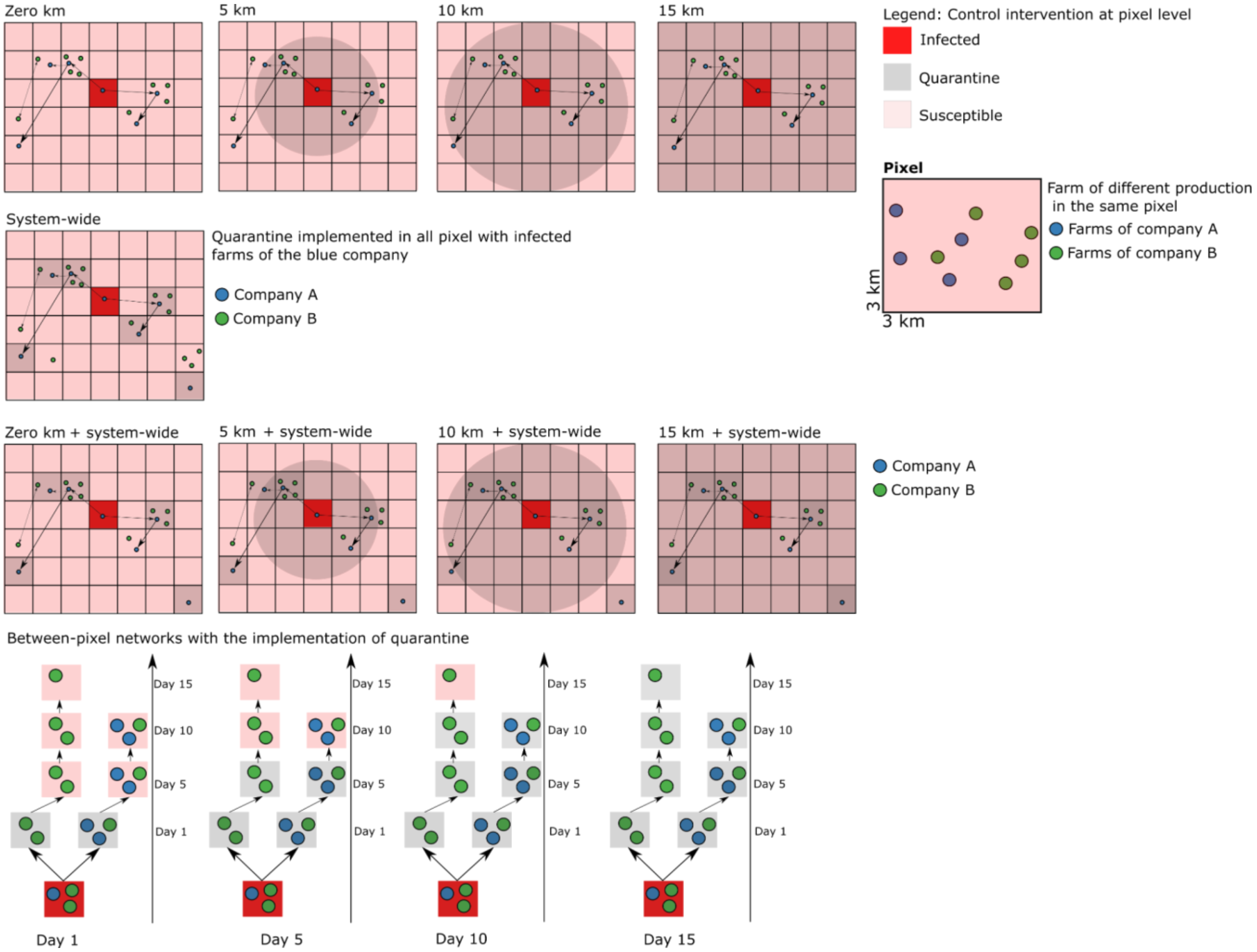


**Figure S2.** Summary of simulated control strategies. The top row (a) shows the intervention deployed by the distance of infected pixels. The mid-row (b) demonstrates how quarantines were deployed to pixels with at least one farm commercial related to an infected pixel, in this scenario all farms of company A are quarantined regardless of recent between-farm movements or distance to where farms may be located. The third row shows the combination of action system-wise and distance-based control. The bottom row demonstrates the network based on backward contact tracing quarantines

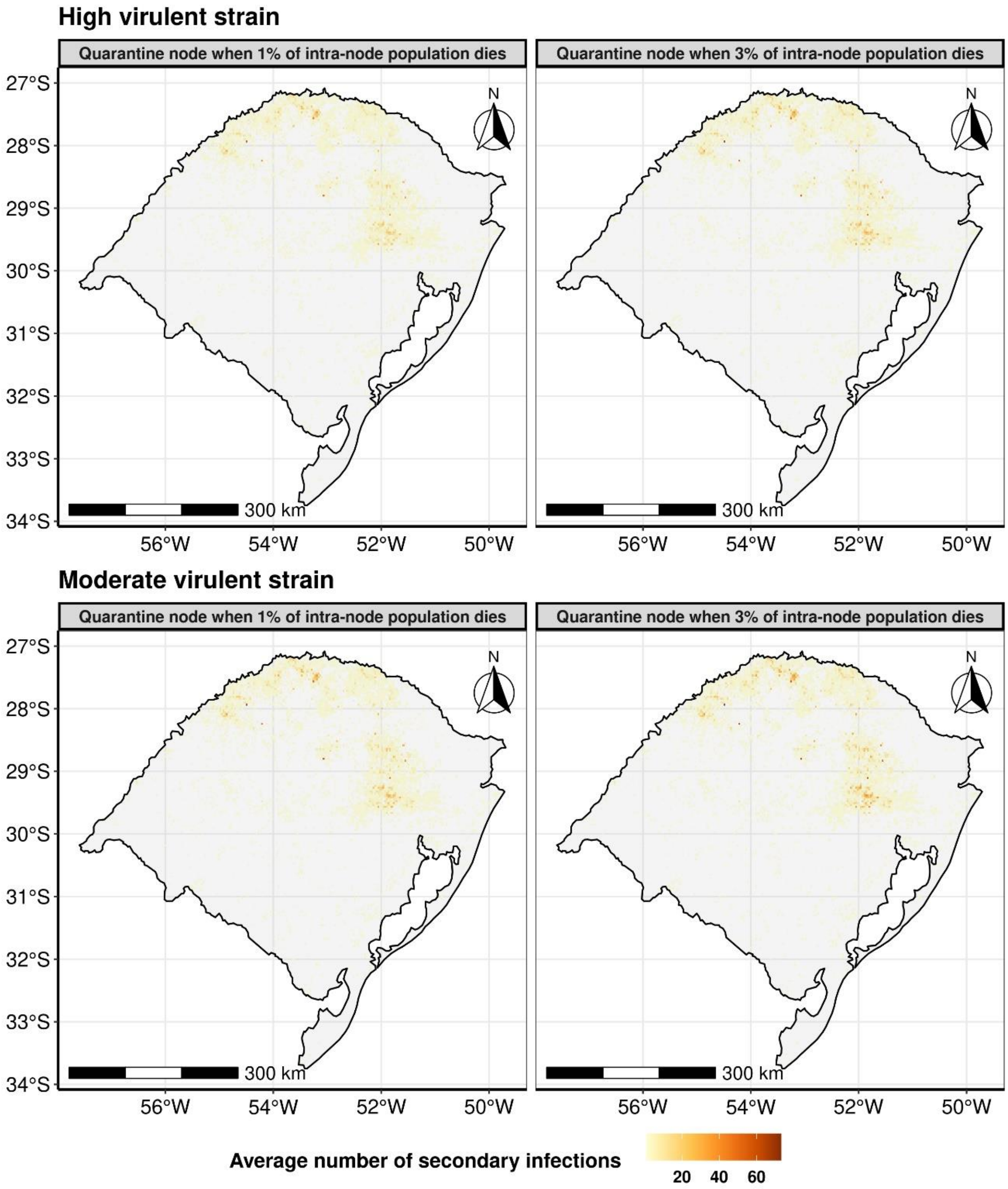


**Figure S3. Average infected pixels (3 km by 3 km).** Pixels highlighted correspond to simulated average secondary cases under the simulated 1% and 3% mortality triggered.

**Table S5.** The proportion of transmission control for all factorial combinations of every variable combination, ASF strains, four mortality triggers, 0%; 1%; 3%; 10%, and control measures.

**Table S6.** The total number of pixels, farms, and pigs is predicted to be quarantine, as a function of ASF strains, control protocols, and mortality triggers. In addition, the column "infection" identifies the simulated infected states as (infected) and uninfected but quarantine as (healthy). Finally the column "Quarantined" divides a binary variable describing if simulations resulted in complete ASF containment or not (i.e. all pixels observed in the Infectious or Exposed state at any point in the simulation pixels transitioned to the Quarantined state at the final timestep.)

**Transmission rate sensitivity analysis:** We carried out on the ρ, which is the probability that an animal transfer from an infected node to a susceptible one will cause the latter to transition to the exposed state, we varied it to be at 50% and 80%. Our results with ρ=100% predicted an average 2 days longer epidemic, an increase in the number of secondarily infected pixels to be 0.2 higher, and spread on average 2km further when compared to 50% and 80% ρ.
